# Characterizing the Resolution and Throughput of the Apollo Direct Electron Detector

**DOI:** 10.1101/2022.09.16.508245

**Authors:** Ruizhi Peng, Xiaofeng Fu, Joshua H. Mendez, Peter S. Randolph, Benjamin E. Bammes, Scott M. Stagg

## Abstract

Advances in electron detection have been essential to the success of high-resolution cryo-EM structure determination. A new generation of direct electron detector called the Apollo, has been developed by Direct Electron. The Apollo uses a novel event-based MAPS detector custom designed for ultra-fast electron counting. We have evaluated this new camera, finding that it delivers high detective quantum efficiency (DQE) and low coincidence loss, enabling high-quality electron counting data acquisition at up to nearly 80 input electrons per pixel per second. We further characterized the performance of Apollo for single particle cryo-EM on real biological samples. Using mouse apoferritin, Apollo yielded better than 1.9 Å resolution reconstructions at all three tested dose rates from a half-day data collection session each. With longer collection time and improved specimen preparation, mouse apoferritin was reconstructed to 1.66 Å resolution. Applied to a more challenging small protein aldolase, we obtained a 2.24 Å resolution reconstruction. The high quality of the map indicates that the Apollo has sufficiently high DQE to reconstruct smaller proteins and complexes with high-fidelity. Our results demonstrate that the Apollo camera performs well across a broad range of dose rates and is capable of capturing high quality data that produce high-resolution reconstructions for large and small single particle samples.

## Introduction

Technological advances have been a major driving factor in the rise of cryogenic electron microscopy (cryo-EM) as a method for determining biological structures [1]. Transmission electron microscopes (TEMs) have had the ability to produce atomic resolution projections since at least the 1980s [2], but until relatively recently, the ability to detect projections of single particles in cryo-EM at high fidelity has been limited. The original detection medium was film, which had multiple limitations, including requiring manual manipulation, the introduction of contaminants into the TEM column, low throughput, and low potential for automation. Contamination, throughput, and automation were dramatically improved by the introduction of charged coupled devices (CCDs) for acquiring TEM images [3–5]. CCD cameras produce electron images indirectly, by first converting electrons to photons using a scintillator, whose optical output is then transferred to a CCD image sensor. The ability to automate cryo-EM data acquisition through digital imaging resulted in a dramatic improvement in throughput, going from 100s to 1000s of images per day [6]. However, digital imaging with CCDs came at the cost of image quality, with both sensitivity and resolution worse than on photographic film [7].

A major breakthrough in the ability to extend the resolution of cryo-EM toward its theoretical limits came with the introduction of direct detectors, including hybrid pixel detectors (HPDs) [8, 9] and monolithic active pixel sensor (MAPS) [10, 11]. For cryo-EM applications, MAPS detectors have become the de facto standard, since HPDs are typically severely limited in pixel array size due to their large pixel size. In either case, because direct detection does not require conversion of the TEM primary electrons to light via a scintillator, the sensitivity and resolution provided by this technology is significantly better than previous detection technologies [12].

There are two types of detection enabled by direct detectors: integration and counting. In integration mode, pixel values represent the total accumulated charge in each pixel. For detectors in which the TEM primary electron passes through the sensor—which includes nearly all direct detectors designed for 200-300 keV electron detection—each TEM primary electron deposits a variable amount of energy in the sensor. This variation is called Landau noise [13, 14] and it reduces the signal-to-noise ratio (SNR) of each image. Additionally, many TEM primary electrons deposit charge in more than one pixel, resulting in a point spread function that attenuates the resolution of each image.

The other type of detection—electron counting—addresses these two deleterious effects [15]. In counting mode, the detection of each TEM primary electron is identified, normalized, and localized, so that all TEM primary electrons are represented by the same intensity and the same shape (typically one pixel). Electron counting significantly improves data quality and has been critical for attaining high-resolution cryo-EM reconstructions [16–19].

While electron counting clearly yields the highest data quality, its primary disadvantage is the requirement for sparse imaging, placing strict conditions on imaging parameters and lowering throughput [17]. The limitations of counting stem from its requirement for TEM primary electrons to be spatially distinguishable in each frame. When the signal from two or more primary electrons overlap, it is impossible to distinguish them as separate events. Thus, a high dose rate on an electron counting camera leads to coincidence loss[16], which has several detrimental consequences. First, since the rate of coincidence loss scales with the exposure rate on the detector, it introduces non-linearity, which may reduce the interpretability of the results. Second, and probably more significant for cryo-EM, coincidence loss causes the detector to miss TEM primary electrons and thus decreases the detective quantum efficiency (DQE) and signal-to-noise ratio (SNR) of the detector [20]. For radiation-sensitive specimens, this is especially problematic since it is not possible to simply use a higher total exposure to compensate for the missed electrons. It is therefore highly desirable to develop large-area electron counting cameras that operate at significantly higher speeds so that coincidence loss is minimized and illumination is not so severely limited.

Increasing counting speed has been challenging for conventional MAPS cameras because these perform electron counting computationally, since these sensors output integrating mode frames consisting of analog pixel values digitized by analog-digital converters (typically 12-bit). Assuming the incident electron beam is sufficiently sparse, these frames are processed in software or firmware to perform thresholding, event identification, localization, and normalization. Alternatively, HPD cameras perform electron counting through complex circuitry within each pixel, which can significantly increase their speed. However, such circuitry requires large pixels, severely limiting sensors to impractically small pixel arrays for cryo-EM [21]. Additionally, current HPDs perform poorly at 200 and 300 kV [22], which are the most popular accelerating voltages used for cryo-EM.

The Apollo (Direct Electron LP, San Diego, CA USA) is a new high-performance direct detector that addresses these challenges by using a novel event-based MAPS detector custom designed for ultra-fast electron counting. This sensor is composed of 4096 × 4096 physical pixels, with 8 µm pixel size. To minimize noise resulting from the pixels not returning to the same baseline after detection, so-called reset noise, each pixel includes on-chip correlated double sampling (CDS) functionality [23], so that noise subtraction occurs automatically within each pixel prior to readout. The upper and lower periphery of the sensor, shielded from the electron beam, include two additional on-chip structures: a sense amplifier and a priority encoder. The sense amplifier performs thresholding and identifies detection events, producing binary output representing either the presence or absence of a detected electron in each pixel. This output is passed to the priority encoder, which packages and digitally outputs groups of pixels containing detection events. Therefore, readout from the Apollo sensor is digital, containing a stream of detection events. This digital output is passed to on-board field-programmable gate arrays (FPGAs), which discriminate and calculate the super-resolution centroid of each separate detection event. Detection events are accumulated in on-board quadruple data rate (QDR) memory over a defined time period, before the dose-fractionated electron-counted super-resolution movie frame is output to the computer for storage and/or display.

Here, we have characterized the Apollo camera in terms of throughput and resolution. We characterized the physical performance of the camera by measuring the noise power spectrum (NPS), modulation transfer function (MTF), and detective quantum efficiency (DQE). These results were extended by testing the performance of the camera for single particle cryo-EM by collecting data on two common test samples, the 481 kDa apoferritin and 149 kDa aldolase.Apoferritin data were collected at three different dose rates to characterize the influence of coincidence loss on single particle throughput and resolution. Finally high-resolution apoferritin and aldolase datasets were collected and reconstructed to 1.66 Å and 2.24 Å respectively. Our results demonstrate that the Apollo camera performs well across a broad range of dose rates and is capable of capturing high quality data that produce high-resolution reconstructions for large and small single particle samples.

## Methods

### Apollo coincidence loss estimate

Apollo was installed on a Titan Krios (Thermo Fisher Scientific, Waltham, MA USA) and was operated at its standard temperature of −25° C. A gain reference was acquired consisting of a cumulative exposure of ∼4000 electrons per physical pixel, with the dose rate set to ∼30 electrons per physical pixel per second (eps). The pixel sizes at nominal magnifications from 14,000× to 120,000× were calibrated using images acquired of a crossed line grating replica (Electron Microscopy Sciences, Hatfield, PA USA), processed with DE_MagCal.ijm macro (available at https://github.com/directelectron/imagej-macros) for Fiji/ImageJ [24].

To assess coincidence loss, gain correction on Apollo was turned off so that raw counts were acquired and saved to disk. The specimen was removed from the column, so that images acquired by Apollo would be uniform flood-beam illumination through vacuum. In nanoprobe mode, the C2 lens was adjusted to 56.319%, which provided uniform illumination over the field-of-view of the camera across a broad range of magnifications and beam brightness settings. Intensity zoom was turned off. With the illumination (C2 and C3 lenses) kept constant, a series of 1 s exposures were acquired on the Apollo, at each magnification ranging from 14,000× to 120,000×. This magnification series was repeated at four different C3 lens settings: 35.229%, 36.524%, 37.188%, and 37.889%. The value of a single detected electron on the Apollo is 16, and there are 4 super-resolution pixels per physical pixel. Therefore, the detected dose rate in electrons per physical pixel per second was calculated for each image by determining the mean pixel intensity for each super-resolution image, then dividing that by 16 to get counts per electron and finally multiplying by 4 to get counts per physical pixel.

The input dose rate was estimated by assuming zero coincidence loss at for the lowest intensity image in each magnification series (that is, the image acquired at the highest magnification) and then scaling that dose rate to the other magnifications based on the calibrated pixel size at each magnification. Results were fit based on least-squares deviation to a previously-published equation for coincidence loss [18], adapted to use units of eps instead of electrons per physical pixel per frame (epf), since the concept of an internal frame rate for counting is not strictly applicable for Apollo. The equation used was:

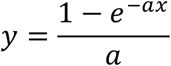

where *y* is the measured (output) dose rate in units of eps, *x* is the input dose rate in units of eps, and *a* is a constant whose value was fit to the experimental data.

The rotationally-averaged power spectrum for each of a similarly acquired series of images was calculated in Fiji/ImageJ by calculating the FFT of a 4096×4096 cropped region from the center of each super-resolution image, using the “Raw power spectrum” option, and then applying the Radial Profile plugin (https://imagej.nih.gov/ij/plugins/radial-profile.html). Each rotationally-averaged power spectrum was normalized by its average value between 1.5× and 2.0× physical Nyquist.

### Apollo performance characterization

With no specimen in the TEM column, the magnification was set to a nominal value of 3800× in microprobe mode. Illumination was set to 54.140% C2 and 37.790% C3, resulting in a detected reading of 12.0 eps on Apollo, which corresponded to an input dose rate of 12.5 (∼4% coincidence loss) based on the coincidence loss curve previously calculated for the camera. The 200 µm selected area aperture was inserted and centered, so that the edges of the beam were visible on the images acquired by Apollo.

The TEM beamstop was inserted and centered on the field-of-view of Apollo, waiting at least 2 minutes after inserted the beamstop to ensure it was no longer moving. A series of 16 acquisitions with 5.2 s exposure time were acquired and then summed in Fiji/ImageJ, resulting in a final image with ∼1040 electrons per physical pixel total. The modulation transfer function (MTF) was calculated using ASI_MTF.ijm (https://github.com/emx77/ASI_MTF) after selecting a straight region from the edge of the beamstop.

Similarly, the TEM beamstop was removed and a series of 16 acquisitions with 5.2 s exposure time were acquired and loaded into Fiji/ImageJ as a stack and cropped to the central 4096×4096 region. DQE(0) was calculated using DE_DQE0.ijm (https://github.com/directelectron/imagej-macros/blob/main/DE_DQE0.ijm), which calculates DQE(0) using the noise binning method [7]. Subsequently, DQE was calculated according to the following equation:

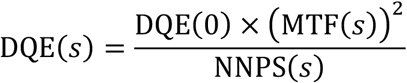

where NNPS is the normalized noise power spectrum, which was set to unity across all spatial frequencies, as is the case in electron counting in the absence of coincidence loss.

### Mouse heavy chain apoferritin

pET24a plasmid with mFTH1 gene was a gift from Dr. Masahide Kikkawa [25]. We amplified the plasmid in DH5α cells and confirmed the sequence by DNA sequencing. The expression strain was obtained by transformation of this plasmid into BL21(DE3). Expression was performed by growing BL21(DE3) in two liters of LB with 50ug/mL kanamycin at 37° C. Finally, IPTG at a final 1 mM concentration was added when OD_600_ reached 0.6. After 3 hours of induction, cells were harvested by centrifugation at 4,000× g for 20 minutes (4° C), resuspended in lysis buffer (30 mM HEPES pH 7.5, 300 mM NaCl, 1 mM MgSO4, freshly added 1mg/mL lysozyme and protease inhibitor cocktail (ThermoFisher Scientific). Resuspended cells were lysed by sonication. Cell debris were centrifuged at 20,000 g for 1h (4° C). The supernatant was incubated at 70° C for 10 mins to denature unwanted proteins. The mixture was centrifuged again at 20,000× g for 1h (4° C) to remove precipitated proteins. Ammonium sulfate at a final concentration of 60% (w/v) was added to the cleared supernatant and stirred on ice for 10 minutes. Precipitated apoferritin was collected by centrifugation at 14,000 g for 30 minutes (4° C), resuspended in 20 mL buffer A (30 mM HEPES pH 7.5, 20 mM NaCl, freshly added 1 mM DTT) and dialyzed against 2×1L buffer A (4° C). Apoferritin solution was loaded onto a HiTrap Q HP 5mL anion exchange column. The column was washed with 5 column volume of buffer A, eluted with a linear gradient of buffer B (30 mM HEPES pH 7.5, 500 mM NaCl, freshly added 1 mM DTT) over 10 column volumes. Elution was confirmed by SDS-PAGE. Fractions containing pure apoferritin were pooled together and concentrated by 100 kDa PES concentrator (Corning) to final 200 μL. Concentrated sample was centrifuged at 15,000 g for 15 minutes (4° C), loaded onto superose 6 increase 10/300 GL column and eluted with buffer C (30 mM HEPES pH 7.5, 150 mM NaCl, freshly added 1 mM DTT). Elution was confirmed by SDS-PAGE. Fractions containing pure apoferritin were pooled together. Concentration was estimated by nanodrop as 8.3 mg/mL. Apoferritin was diluted to 4mg/mL before flash-freezing in liquid nitrogen for long-term storage.

### Rabbit muscle aldolase

Rabbit muscle aldolase vitrified on an UltrAuFoil R1.2/1.3 300-mesh grid was kindly provided by Dr. Gabriel Lander [26]

### Cryo-EM grid preparation

Two different batches of apoferritin grids were made. For each a 4 mg/mL apoferritin aliquot was thawed on ice and centrifuged at 15,000 g for 20 minutes to remove aggregates. UltraAuFoil R1.2/1.3 300 Mesh grids were plasma cleaned for 50 seconds for the first batch and 20 seconds for the second batch at 7 Watts with 25% oxygen, 75% argon (Gatan Solarus 950). Cleaned grids were loaded onto Vitrobot Mark IV at 100% humidity (4° C). 4 μL apoferritin was applied to the grid. For the first batch, the grids were blotted for 1 second at 0 force, and for the second batch, the grid was blotted for 4 seconds at 0 force before being vitrified in liquid ethane.

### Cryo-EM data acquisition

All data, unless indicated otherwise, were collected on a Titan Krios microscope (ThermoFisher Scientific) operated at 300 kV with a 50 μm C2 aperture and 100 μm objective aperture and equipped with a DE Apollo direct detector. The electron beam in nanoprobe mode was aligned through Direct Alignment in the Titan GUI. Coma-free alignment was performed by acquiring a Zemlin tableau in Leginon [27, 28]. Cryo-EM images were acquired using the Leginon software and pre-processed using the Appion package[29]. Apollo camera was set in 8k × 8k super-resolution mode. For both apoferritin and aldolase, movies were collected at a magnification of 75000× with the corrected pixel size of 0.599 Å. The movie frame rate was fixed at 60 frames per second for all datasets.

Aldolase movies were collected with a random defocus range of −0.8 μm to −1.5 μm. The movies were collected at a detected dose rate of 15 e^−^/pixel/second (eps)—corresponding to ∼16 eps input dose rate after accounting for coincidence loss—for 1347ms, resulting in a total exposure of around 60 e^−^ / Å^2^. Movies were binned 2× during motion correction.

Apoferritin movies were collected with a random defocus range of −0.4 μm to −1.0 μm. Data collection were done in two sessions. In the first session, movies were collected at three detected dose rates respectively, 15 eps, 30 eps, and 60 eps corresponding to input dose rates of 16 eps, 34 eps, and 78 eps, respectively. For all three dose rates, the exposure time was set such that the total dose per exposure was ∼ 60 e^−^ / Å^2^. In the second session the dose rate was kept constant at 15 detected eps. Movies were binned 2× during motion correction

### Aldolase cryo-EM data processing

A total of 8682 movies were imported into cryoSPARC 3.3.2 [30], and patch motion corrected in 6×6 segments. CTF was estimated by CTFFIND4 [31]. 5557 micrographs with estimated defocus from −1.5 μm to −0.4 μm, and maximum resolution higher than 5 Å were selected for further processing. EMD-21023 was used to generate 2D templates which were low-pass filtered to 15 Å and used for template-based particle picking. A total of 3,805,094 particles were picked and boxed into a 512-pixel box. Two rounds of 2D classifications were done using a mask of 130 Å diameter, 2D averages with clear high-resolution features were selected, resulting in a set of 1,824,301 particles. 722,778 particles from the best class of 3D heterogeneous refinement were selected for further processing. The first homogeneous refinement resulted in a 2.55 Å map. CTF refinement was performed by first refining magnification anisotropy, then refining beam tilt, spherical aberrations; and finally refining per-particle defocus. Non-uniform refinement yielded a resolution of 2.31 Å. Bayesian polishing and another round of CTF refinement resulted in a 2.24 Å map.

### Apoferritin cryo-EM data processing

For the highest resolution dataset, a total of 3804 movies were imported into Relion3.1 [32], motion was corrected in 5×5 segments with a B-factor of 150 by MotionCor2 [33], and CTF was estimated by CTFFIND4. 2050 micrographs with estimated defocus from −1.3 μm to −0.4 μm and CTF maximum resolution higher than 5 Å were selected for further processing. A previously reconstructed 2.6 Å apoferritin map was used to generate 2D templates which were low-pass filtered to 10 Å and used for template-based particle picking. 401,410 particles were picked in a 480-pixel box. Particles were 2x binned into 240-pixel boxes for classification. 2D averages with clear features were selected, resulting in a set of 312,431 particles. Particles were 3D classified in four classes for 25 iterations. The class with the highest resolution was selected for reconstruction, resulting in a set of 286,970 particles. 3D auto-refinement yield a 2.46 Å resolution map. Briefly, two rounds of stepwise CTF refinement consisting of beam tilt refinement, anisotropic magnification refinement, and local CTF parameters refinement were done between multiple rounds of 3D auto-refinement. This cycle was repeated until there was no improvement in map resolution. Particles were then reconstructed with Ewald sphere correction. These yielded a final 1.66 Å resolution map.

Datasets collected with different dose rates were processed in the same manner as above. For the 15 detected eps dataset, 972 movies were collected resulting in 65,516 particles after classification and a 1.68 Å map. For the 30 detected eps dataset, 1478 movies were collected resulting in 93,866 particles after classification and a 1.68 Å map. For the 60 detected eps dataset, 869 movies were collected resulting in 94,346 particles after classification and a 1.87 Å map.

### Apoferritin modelling

Mouse heavy chain apoferritin was remodeled with using PDB 7A4M [18] as the initial model. Two apoferritin maps were imported into PHENIX [34]. The asymmetric units were extracted from the EM densities and the initial model (stripped of ions and waters) was fit into the resulting maps. Waters and ions were added and the fit of individual residues in the maps were improved in COOT [35]. The structures then underwent refinement and symmetrization in PHENIX.

### Aldolase modelling

Rabbit muscle aldolase was remodeled using PDB 7KA4 [36] as an initial model. The aldolase map was imported into PHENIX [34] which was used to extract the asymmetric unit from the EM density. A single chain from 7KA4 and the asymmetric EM density were refined in ISOLDE [37] remodeling side chains to match the density. After the structure was refined it was imported back into PHENIX, symmetrized, and refined to complete the structure.

## Results

### Characterization of Apollo imaging performance

We first characterized the performance of the Apollo across a range of dose rates to assess the linearity of the camera and the rate of coincidence loss. Coincidence loss has the effect of reducing the linearity [16] of detected electrons compared to incident electrons, which not only reduces the overall detection efficiency, but also dampens the low frequencies of the noise power spectrum [20]. We assessed coincidence loss by examining both effects.

After normalizing the noise power spectrum for each dose rate to unity at super-resolution Nyquist, the dampening of the low spatial frequencies in the noise power spectrum due to coincidence loss was evident (Fig 1A). As expected, the amount of dampening increased with increasing dose rate, as coincidence loss was more severe at high dose rates than low dose rates. We calculate the mean value of the noise power spectrum across spatial frequencies from 0.06 to 0.08 physical Nyquist frequency, which appeared to be the approximate location of the minimum in each noise power spectrum curve. It has been previously shown that the minimum value in the power spectrum at low spatial frequencies corresponds to the amount of coincidence loss in the image [16]. However, because of the noise present in the rotationally averaged power spectrum curves, we chose not to rely on the power spectrum measurements to precisely quantify the coincidence loss rate on the Apollo. Nevertheless, the power spectrum curves imply that coincidence loss is small at 15 eps and negligible at lower dose rates, since the corresponding power spectrum curves showed minimal dampening.

**Fig. 1.**
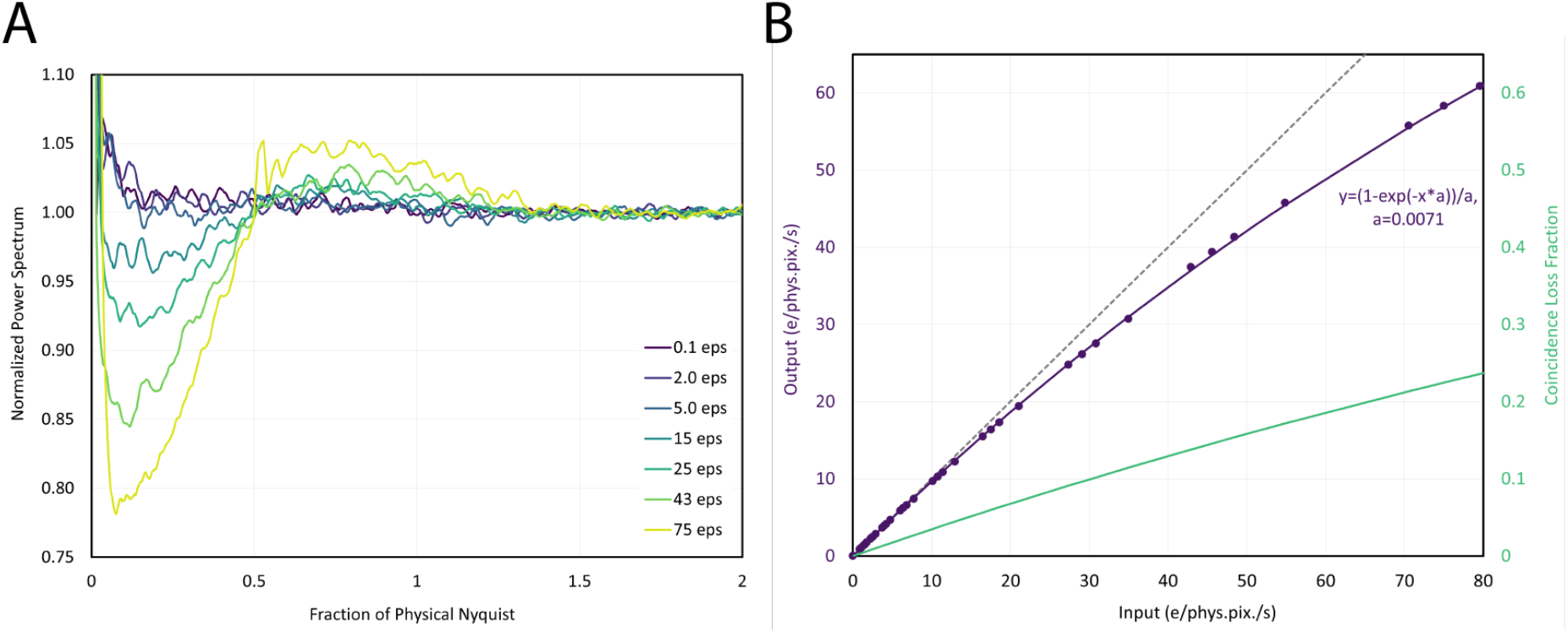
Coincidence loss for the DE Apollo. A) Noise power spectra for different input dose rates. Suppression of low frequency power can be observed with increasing dose rate. B) Dose response curve. The dashed line represents the dose response condition in the absence of coincidence loss. The solid line represents the fit to the measured detected dose rate with increasing the input dose rate (blue circles). The green line is the coincidence loss fraction.

Notably, the noise power spectrum was unity even at an ultra-low dose rate of 0.1 eps, where detector noise would be expected to have a large effect, if present. At the sensor threshold value used in our experiments, we detected a false positive (noise) rate of 2.86 × 10^−6^ events per pixel per second, based on acquisition of images with the TEM beam valve closed. Even at a low dose rate of 0.1 eps, this false positive rate is negligible (<0.003%).

To quantify the coincidence loss on the Apollo, the detected dose rate on the camera was plotted against the expected (input) dose rate (Fig. 1B). In the absence of coincidence loss, this plot should be perfectly linear. But as the dose rate outpaces the rate of electron detection, the plot will deviate from linearity by precisely the amount of coincidence loss at each rate.

The dose rate response curve for Apollo fit well with a previously published equation for coincidence loss [18], with the units changed from dose rate per frame to dose rate per second. For cameras that output integrating-mode frames to be counted through post-processing, using units based on the frame rate is sensible. However, because Apollo performs electron counting in real-time through on-chip and on-board electronics, the concept of counting frame rate is no longer applicable. Instead, camera performance and coincidence loss must be evaluated on the basis of dose rate per unit time. Conveniently, all other cameras, regardless of their counting strategy, can be evaluated on the basis of these units. Therefore, coincidence loss versus eps provides an effective metric for comparing the performance of various electron counting cameras.

We found that the coincidence loss at 300 kV was ∼5% at an input dose rate of 15 eps, ∼10% at an input dose rate of 30 eps, and ∼20% at an input dose rate of 66 eps (Fig. 1B). The estimated coincidence loss based on power spectrum dampening approximately matched our quantitative assessment of coincidence loss based on input and output dose rates (Fig. 1B).

Increasing the input dose rate beyond ∼80 eps (corresponding to a detected dose rate of ∼60 eps) resulted in saturation, with strong gradients across each segment in the image, likely due to the count rate exceeding the bandwidth of the FPGAs and on-board memory. These artifacts were not present at lower dose rates.

For the remainder of the manuscript, we will explicitly distinguish dose rate measurements by denoting whether they are the measured detected dose rates or the input dose rates as estimated from the dose response curve in Fig. 1B.

The resolution and signal-to-noise ratio (SNR) of a detector can be estimated by the modulation transfer function (MTF) and detective quantum efficiency (DQE). Because these measurements—especially DQE—can depend heavily on the imaging conditions used [20, 38], they are most relevant if they are acquired under experimental conditions similar to those that would be used in practice for cryo-EM data acquisition. We chose to perform these calculations at 12 detected eps, which is realistic for high-resolution cryo-EM data acquisition. At a magnification yielding 0.6 Å/physical pixel sampling, this corresponds to a dose rate on the specimen of 33 e^-^/Å^2^, which would enable single-particle data acquisition with exposure time in the range of 1-2 seconds, depending on the desired total exposure.

The MTF was estimated by acquiring images of the TEM beamstop, although this tends to result in an underestimate of the MTF due to the large distance between the beamstop and the sensor, the rounded shape of the beamstop, and the roughness of the beamstop edges. We tried to minimize the latter effect by selecting a region-of-interest that appeared relatively straight by visual inspection (Fig. 2A). Based on this region, the MTF was approximately 0.84 at half-Nyquist and 0.5 at Nyquist (Fig. 2B). Using the noise binning method [7], DQE(0) was calculated as 0.937, resulting in a DQE of 0.66 at half-Nyquist and 0.23 at Nyquist (Fig. 2B).

**Fig. 2.**
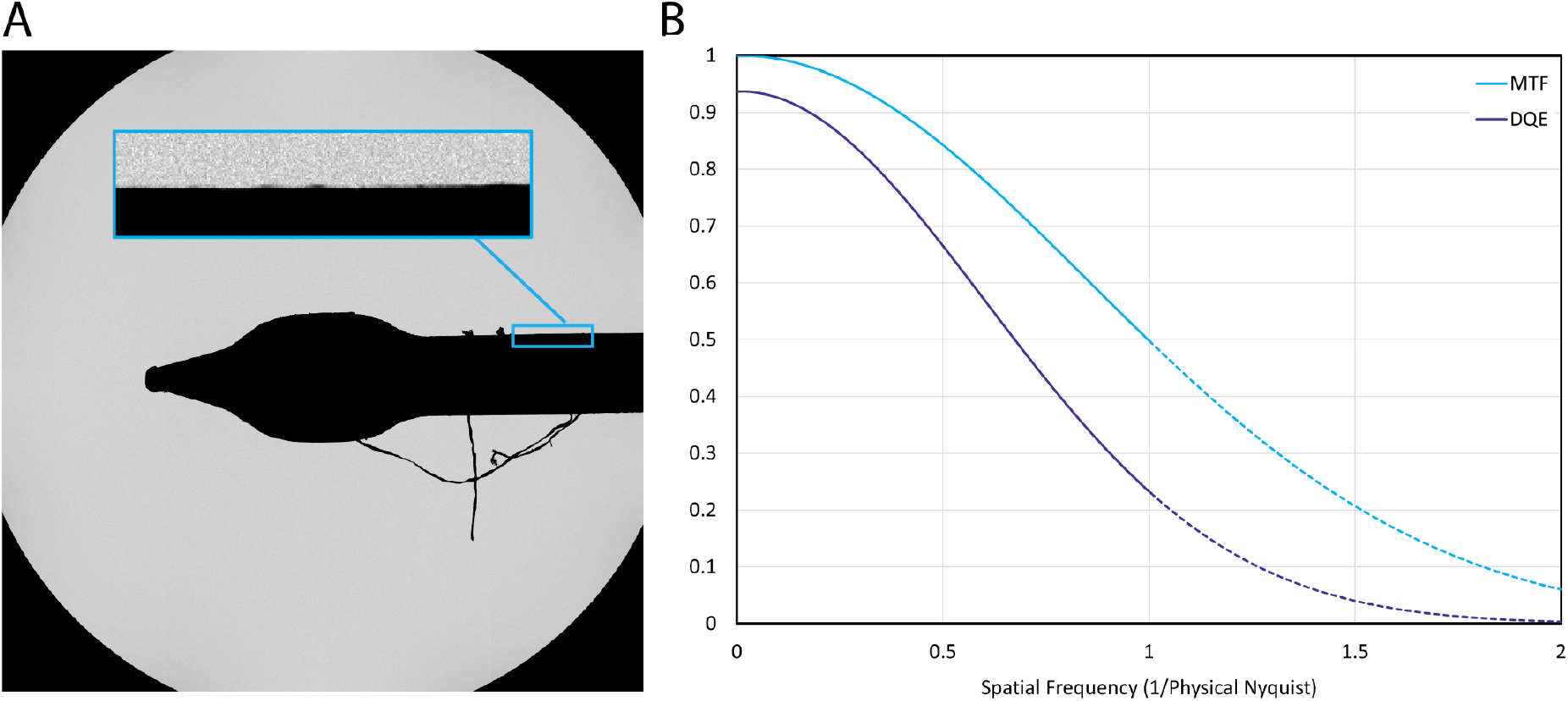
Calculating the MTF and DQE. A) Image of the beam stop on the Titan Krios. The inset shows a straight portion of the beamstop used to calculate the MTF. B) MTF (cyan) and DQE (purple) plots for the Apollo

The coincidence loss at 12 detected eps is ∼4% and the normalized noise power spectrum (NNPS) at this exposure rate shows little dampening at low spatial frequencies, with the entire NNPS fluctuating around unity across all spatial frequencies. However, to avoid adding irrelevant noise to the DQE curve and to artificially inflating the DQE at low spatial frequencies due to the small amount of dampening in the NNPS, we treated the NNPS as exactly unity at all spatial frequencies.

The resulting DQE curve was approximately 0.66 at half-Nyquist and 0.23 at Nyquist (Fig. 2B). DQE remained above 0.10 for super-resolution spatial frequencies, up to ∼1.25× physical Nyquist.

### Characterizing the performance of the Apollo on cryo-EM data

The Apollo was implemented in Leginon [28] to facilitate automated data collection. Single particle apoferritin cryo-EM data were collected at 15, 30, and 60 detected eps (corresponding to 16, 34, and 78 input eps, respectively) to characterize the performance of the camera at different dose rates. The data were all collected on the same grid and processed in the same way. The total dose for each exposure was 60 e^-^/Å^2^ for each dose rate, and that resulted in exposure times of 1264, 666, and 333 ms respectively. Given the fixed movie frame rate, there was necessarily a difference in the number of movie frames for each condition, with 76, 40, and 20 frames respectively. It is important to note that this difference could affect the ability to correct for beam induced motion and beam damage for the different datasets. Greater than 850 micrographs were collected at each condition resulting in 183,110 particles for the 15 detected eps data, 251,816 for 30 eps, and 203,117 particles for 60 eps. These were processed identically in Relion [39] and resulted in reconstructions at 1.68 Å, 1.68 Å, and 1.87 Å respectively (Fig. 3). All three maps had the expected features for high-resolution maps including holes in ring structures, though the 60 eps 1.87 Å map had holes only in six membered rings while the other two maps had resolvable holes in both six and five membered rings (Fig. 3B).

**Fig. 3.**
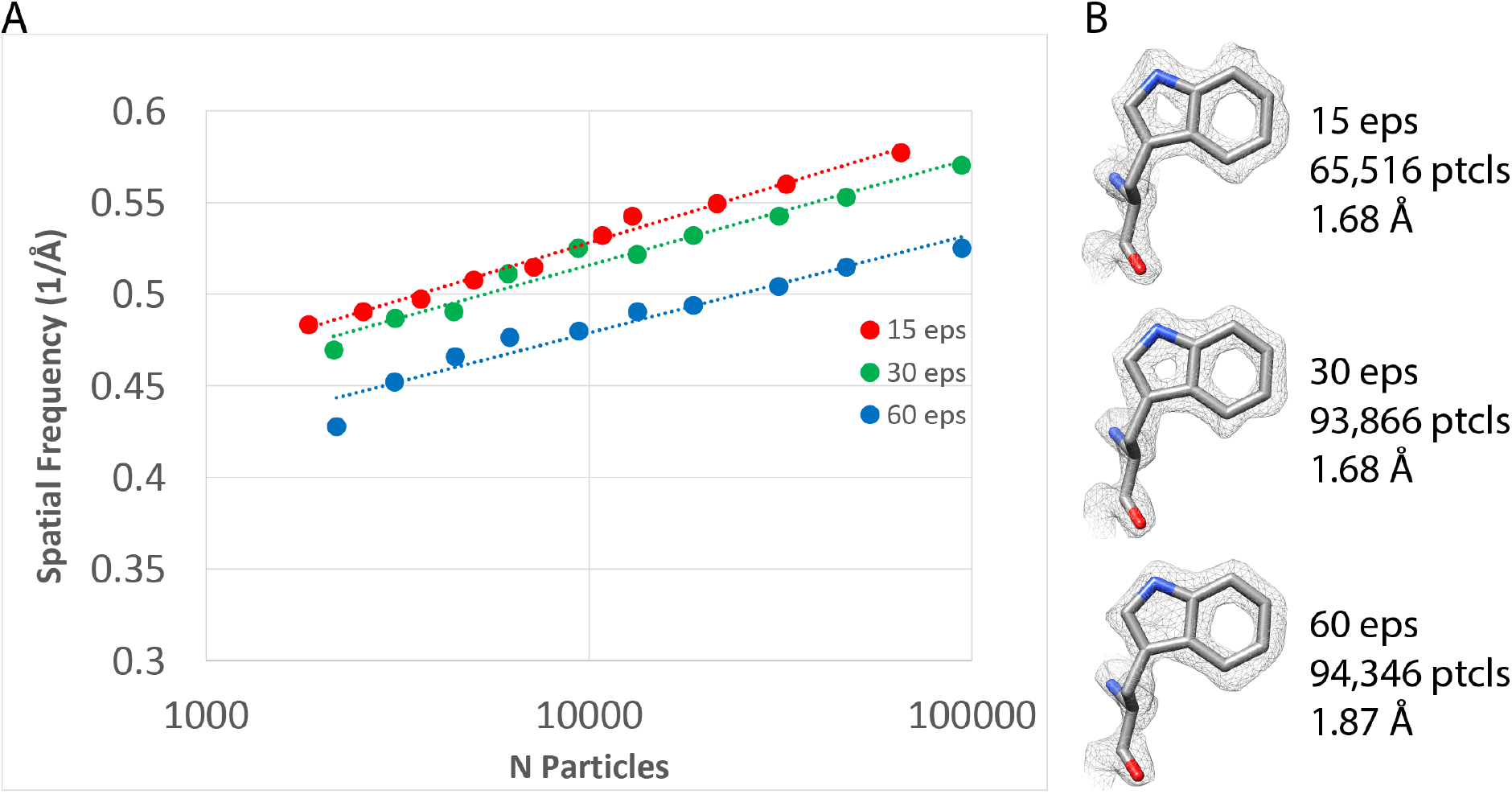
Dose rate comparison. A) ResLog plots of apoferritin collected at 15, 30, and 60 detected eps. B) Tryptophan density for apoferritin collected at 15, 30, and 60 detected eps.

We have previously shown that plots of resolution versus the logarithm of the number of particles (ResLog plots) are linear and the ResLog slopes correspond to overall image quality while ResLog intercepts (the extrapolated resolution from only 1 particle) correspond to the average accuracy of particle alignment and classification [40]. We made ResLog plots for the 15, 30, and 60 detected eps data sets (Fig. 3A). The ResLog slopes were similar for each dataset, but the trend showed that the slope correlated with dose rate. The slopes were 0.063, 0.058 (∼8% worse), and 0.054 (∼15% worse), for the 15, 30, and 60 eps data, respectively. This indicates that correctly aligned particles from data with lower coincidence loss contributed to a greater increase in overall map resolution compared to data with higher coincidence loss. In other words, the information content was greater when coincidence loss was reduced. Similarly, the ResLog intercepts for the 15 and 30 detected eps data were nearly the same while the 60 eps data was substantially lower (worse), implying that coincidence loss affects the average accuracy of particle alignment. Since the low frequencies are critical for single particle alignment, we believe this is likely due to the negative effect that coincidence loss has on the the lower frequencies in the DQE

### High-resolution reconstruction of apoferritin

It was observed that images for the apoferritin grid in the dose rate experiment (Fig. 3) had less than optimal ice, and most images had indications of a water ring in their Fourier transforms. On one hand, the thicker ice emphasizes the importance of DQE for the different dose-rate conditions, but at the same time, it has the potential to limit the resolution of the resulting reconstructions. Thus, another dataset was collected with an ice-optimized apoferritin grid. 3804 images were collected on this grid at a dose rate of 15 detected eps. This resulted in 286,970 particles after classification that reconstructed to 1.66 Å resolution (Fig. 4A). The expected indications of high resolution were present in the reconstruction including holes in 6 and 5 membered rings, the observation of numerous water molecules and ions and the beginnings of indications of hydrogens (Fig. 4B). Despite being collected on a presumably better-quality sample and having more particles, the ice-optimized grid was only 0.02 Å higher resolution than the previous 15 eps map. It is not clear why the resolution is limited to 1.66 Å. The 100 μm TEM objective aperture was inserted during data collection, which could have imposed a resolution cutoff. Another possible resolution limiting factor is that the FSU Titan Krios is equipped with an SFEG instead of the XFEGs or CFEGs that more contemporary TEMs are currently equipped with. The SFEG has more energy spread than the other electron sources which contributes to an uncorrectable chromatic aberration.

**Fig. 4.**
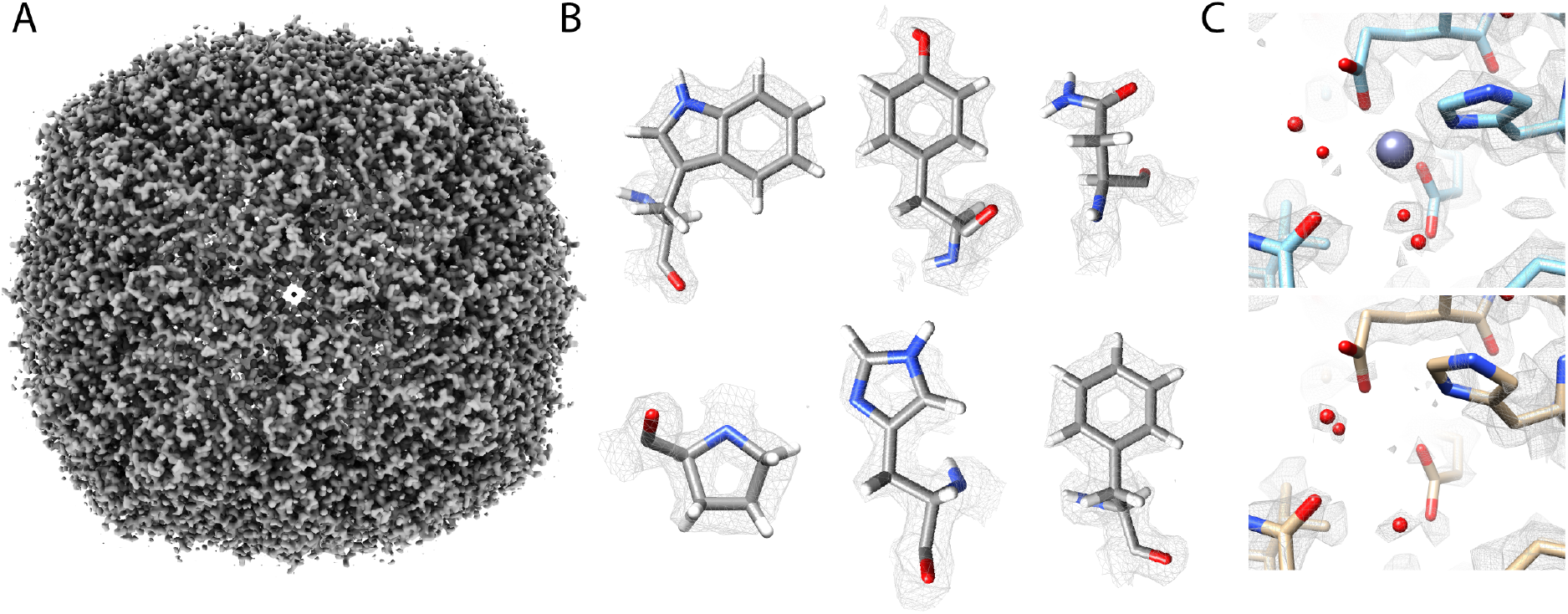
High resolution apoferritin reconstruction. A) Apoferritin density at 1.66 Å. B) High resolution side chain densities with evidence of H densities. C) Zn^2+^ interactions. The top panel shows the apoferritin map with Zn_2+_ density, while the bottom panel shows without. Multiple waters are seen to change positions, as well as a conformational change for His61.

An atomic model was built into the apoferritin density, and the ice-optimized map was compared against the earlier 15 detected eps map. This revealed a surprising difference between the two maps. There is a presumed Zn^2+^ atom chelated between Glu23, Glu58, and His61. The ice-optimized map had clear evidence of the Zn^2+^ atom, similar to other high resolution cryo-EM apoferritin reconstructions [18, 41], whereas it was absent in the earlier 15 detected eps map (Fig. 4C). The resolutions of the two maps were nearly identical at 1.66 Å and 1.68 Å respectively, and they have almost identical features. The two samples were vitrified from the same apoferritin preparation, and the only difference between the two is that one was plasma cleaned for 20 s with 4 s blotting while the other was plasma cleaned for 50 s with 1 s blotting. In addition to the differences in the Zn^2+^ density, there were differences in the water molecules surrounding the Zn^2+^ and the density for chelating residues Glu23, Glu58, and His61 became more well defined indicating that they are better stabilized in the presence of the ion. In the presence of the ion, residues Glu23, Glu58, and His61 are stabilized and two waters fill out two of the three the remaining ligand positions leaving one remaining. In the absence of the ion, one of the liganded waters moves 0.5 Å away from the Zn^2+^ binding site, and the other liganded water is no longer present in the binding site. These lead to changes in the positions of multiple surrounding waters. Finally, the imidazole ring His61 rotates 22 ° away from the Zn^2+^ binding site suggesting an induced fit mechanism for the ion binding. There are no other observable changes in the peptide backbone. These data indicate that there is a network of interactions between the Zn^2+^ ion, surrounding waters, and side chains that rearrange based on the presence and absence of the ion.

### High-resolution reconstruction of aldolase

In order to see how the camera performed on a more challenging specimen, single particle data were collected on rabbit muscle aldolase, which is a 149 kDa tetramer with D2 symmetry.

722,778 particles of aldolase were collected and processed in cryoSparc. This resulted in a reconstruction at 2.24 Å resolution (Fig. 5A). The map showed high quality features including clear side chain densities and some high-resolution features such as holes in the middle of six membered rings (Fig. 5B). The high quality of the map indicates that the Apollo has sufficiently high DQE to reconstruct smaller proteins and complexes with high-fidelity.

**Fig. 5.**
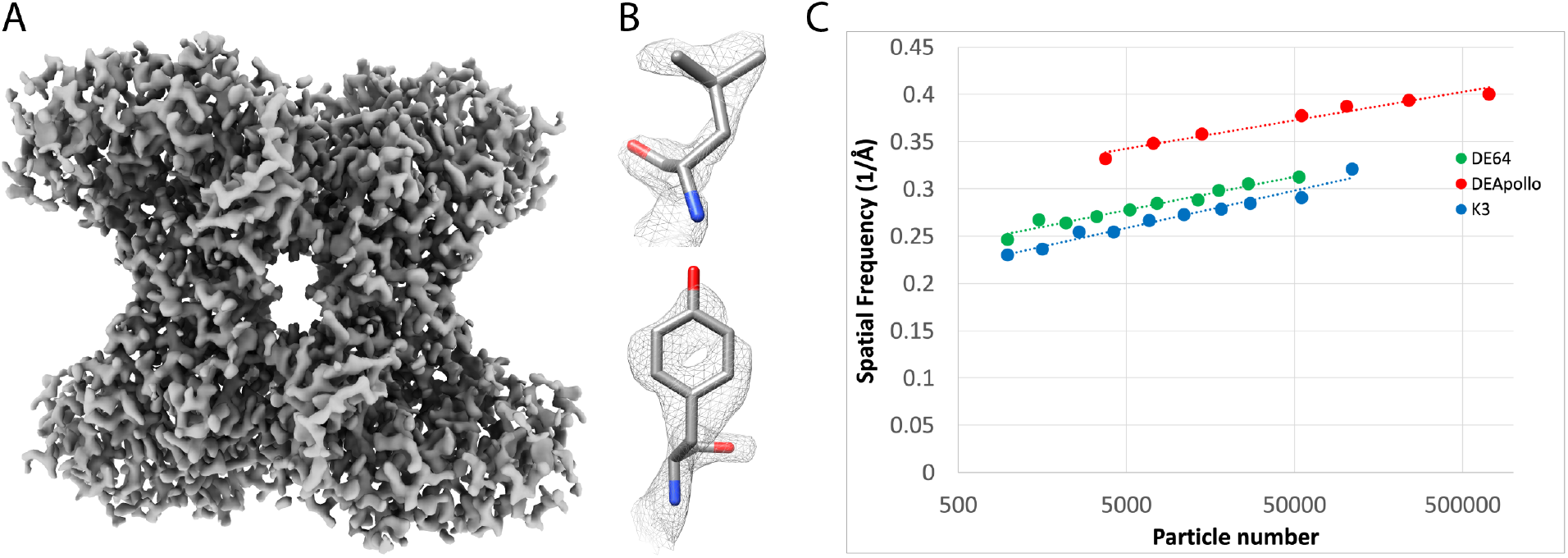
High-resolution aldolase reconstruction. A) 2.24 Å aldolase reconstruction. B) High-resolution details in the side chain densities. C) ResLog plots of aldolase data collected on a DE Apollo (red), DE-64 (green), K3 (blue).

We also compared the performance of the Apollo against the DE-64 (Direct Electron) and K3 cameras (Gatan, Pleasanton, CA USA) using aldolase grids. Aldolase datasets were collected with each of the cameras. The K3 and DE-64 data were collected on the same grid while the Apollo grid came from a different grid in the same preparation session. 59,636 particles were collected for the DE-64 and 110,165 for the K3, and they reconstructed to 2.94Å, and 2.90 Å respectively. ResLog plots were made for each of the refinements (Fig. 5C). As we demonstrated on the apoferritin sample, there can be substantial variation in the sample from grid to grid so only the DE-64 and K3 data can be unambiguously compared. The ResLog slopes were nearly the same for each of the datasets indicating that well aligned particles contribute similarly to the reconstruction for each of the cameras. The ResLog intercepts were close for the two cameras with the DE-64 having a slightly higher intercept. This indicates that the particles from the DE-64 were slightly more alignable than the K3, however, the throughput on the DE-64 is slower than the K3 with an image-to-image time of 1.2 min/image for the DE-64 compared to 14.3 s/image with the K3. Thus, the lower imaging performance of the K3 was made up for by an increase in acquisition speed. The Apollo data had a substantially higher ResLog intercept than the other two cameras. The data were collected from a different grid, so they are not directly comparable, but by visual inspection, the grids appeared to be of similar quality judging by ice thickness and monodispersity of particles. Nonetheless, the reconstruction went to substantially higher resolution than the other cameras, and the time to return an 8k x 8k super resolution image is around 3 seconds compared to the DE-64 which is on the order of a minute.

### Throughput on the Apollo

Data collection throughput in cryo-EM is highly dependent on the data collection strategy which, in turn, depends on the pattern of holes on the EM grid, how frequently one focuses, how many exposures can be collected per hole, etc. In order to assess the acquisition speed of the Apollo independently of those factors, we used Leginon scripting to take back-to-back images on the Apollo. The Apollo has a fixed output movie frame rate of 16.6 ms/frame to the computer, so the number of movie frames for a given exposure is linearly dependent on the exposure time. A consequence of this is that the time it takes to write out the movie files increases with exposure time so there is additional processing overhead as exposure time goes up. In our tests on taking and saving super-resolution 8192 × 8192 pixel movies in 8-bit MRC format, a 1.264 s exposure (the same as our 15 detected eps apoferritin dataset) completed in 3.595s (2.3 s overhead), a 0.666 s exposure completed in 2.303 s (1.6 s overhead), and a 0.333 s exposure completed in 1.542 s (1.2 s overhead). On our system, there was an additional 9.3 s per image overhead in Leginon to transfer the 8k images over the network and process them, but it is expected that this can be practically eliminated with better networking and a better data collection computer.

## Discussion

An important aspect of direct electron detection cameras is that they record data well across all spatial frequencies. The quality of the images for a detector is quantified using the detective quantum efficiency (DQE), which is defined as DQE(ω)=SNR_out_^2^(ω)/SNR_in2_(ω). The highest quality imaging occurs at frequencies where the DQE approaches 1, and the theoretical maximum for a detector with square pixels have DQE(0)=1 and DQE(Nyquist)=0.4. Due to the elimination of Landau noise and the increased precision in localizing each detected electron, electron counting pushes the DQE closer to these theoretical limits, particularly at low spatial frequencies. The detection of low frequency information is particularly important for single particle cryo-EM because the low frequencies are what drives the accuracy of individual particle alignment and classification. For this reason, counting detectors have come to dominate the field for cryo-EM electron detection. However, electron counting requires sparse events distributed across each camera frame and thus it imposes strict limits on the dose rate that may be used for data collection. When the dose rate increases, the per-frame sparsity decreases and it is more likely that multiple electrons will overlap within a single frame and thus be missed, resulting in coincidence loss.

Data collection throughput is another important factor for driving resolution in single particle cryo-EM. There is a logarithmic relationship between resolution and the number of particle images required to achieve it emphasizing the importance of collecting data as rapidly as possible [40, 42]. Throughput with counting detectors, however, is limited by how quickly the camera can count individual electron hits on each individual pixel. Thus, there is an interplay between minimizing coincidence loss by using low dose rates with a concomitant increase in exposure time and maximizing throughput by having the smallest possible exposure time.

The DE Apollo is a new generation of counting direct electron detector that has the potential to increase throughput and resolution for cryo-EM structure determination. The Apollo is based on new technology to perform electron counting in hardware, using an event-based direct detection sensor. This enables the Apollo to perform electron counting on-the-fly at significantly higher speed without sacrificing image quality. We found that coincidence loss is ∼5% at 16 input eps, ∼10% at 34 input eps, and ∼24% at its maximum intensity of 78 input eps. The false positive rate remained very low at 2.86 × 10^−6^ events per pixel per second, likely due to on-chip correlated double sampling (CDS), on-chip thresholding, and digital readout from the sensor. This yielded high DQE, with data quality demonstrated by high-resolution cryo-EM reconstructions. As a demonstration of the data quality and speed of the camera, we achieved sub-2-Å resolution even at a very high dose rate of 78 input eps.

The strong camera performance over a broad range of dose rates provides users with flexibility in their data collection strategy. For projects targeting the highest resolutions, a lower dose rate should be used (e.g., 15 detected eps) in order to minimize coincidence loss and maximize the low frequency DQE. Even the relatively small differences in the ResLog slopes and intercepts that we observed for different dose rates correspond to dramatic differences in the number of particles necessary to reach high-resolution. For example, based on our ResLog plots for apoferritin, increasing the illumination to 30 detected eps instead of 15 detected eps would increase the number of particles needed to reach 1.5 Å resolution by a factor of ∼2.5×. Further increasing the illumination to 60 detected would increase the number of particles needed to reach 1.5 Å resolution by a factor of ∼20×. In either case, even if data acquisition had zero overhead so that the number of particles acquired per unit time scaled perfectly with the exposure time per acquisition, it would be more efficient to acquire data with lower coincidence loss when targeting such high resolution. The recommendation to acquire less data with higher quality is further emphasized by the additional computational demand for image processing and 3D reconstruction required by larger data sets.

There are situations where users may want to sacrifice coincidence loss for throughput. For instance, in the case of specimen screening, there is a need to collect images as quickly as possible to assess sample quality before the go no-go decision can be made for data collection. At the same time, the images must be high enough quality to make informed decisions about sample quality. In these cases, the 60 detected eps imaging condition may be an ideal choice as it affords a 2.3× increase in image acquisition throughput compared to 15 detected eps, and still enables better than 2 Å reconstructions. Similarly for projects with samples with a preferred orientation, rare views, or large compositional heterogeneity that require the maximum possible throughput, it may be advantageous to use the higher dose rate imaging conditions. For example, for 4 Å resolution, increasing the illumination to 60 detected eps from 15 would only require ∼1.4× more particles, which is compensated by the shorter exposure time and thus increased number of particles acquired per unit time.

Our highest resolution reconstruction was of apoferritin at 1.66 Å resolution. The Titan Krios, the data were collected on is a first-generation Titan with an SFEG and without fringe-free data collection. These likely limited the highest possible resolution by chromatic aberration due to beam incoherencies at the highest resolutions and by limitations in throughput by not maximizing the number of images than can be acquired per hole.

The fast imaging speed on the Apollo provides the potential to increase the throughput of cryo-EM data collection. With a 1264 ms exposure, 15 detected eps, and 60 e-/Å^2^, the camera can take images every 3.595 s. We demonstrated that apoferritin collected under these conditions achieved 1.68 Å resolution. There are 86,400 s in a day, and if there was no additional overhead, we could collect 24,033 images per day in this regime. Only 972 images were required at this dose rate to achieve that resolution, so all other things being equal, 24 datasets could be acquired in that time. Of course, this does not include the overhead time it takes to change samples and the myriad activities microscopists do to find “good” regions of a grid. Decreasing the image to image overhead in the data collection software represents a good target for future development in order to achieve this throughput.

The differences in the Zn^2+^ ions, waters, and side chains between different apoferritin datasets was an unexpected observation that was made possible by the high-resolution reconstructions enabled by the Apollo. Surprisingly, despite coming from the same protein purification, one apoferritin grid had clear Zn^2+^ ions while the other had none. This revealed a network of interactions of waters and amino acid side chains that rearrange in the presence and absence of the Zn^2+^ without associated changes in the peptide backbone. It is unknown why these differences arose, but it is likely due to some slight variations in the blotting during plunge freezing. The only other appreciable difference in the two datasets is that the one without Zn^2+^ also had an apparent water ring in the FFTs while the one with Zn^2+^ ions had no evidence of the water ring. Thus, it is possible that the ice thickness during blotting has some effect on the presence and absence of Zn^2+^. It is possible that the thinner ice with the latter sample was due to additional water evaporation during blotting with concomitant increase in Zn^2+^ concentration, but this remains to be proven. Nonetheless, this result highlights the fact that subtle differences in vitrification can have significant differences on the states of cryogenically prepared samples.

## Acknowledgements

Funding for this research was provided by National Institutes of Health grants GM143805 and GM119032 to SMS, and the Apollo was acquired by a supplement to GM139616 to Kenneth Taylor. Maps and models were deposited to the Protein Data Bank and EM Data Resource: TBD, TBD – apoferritin with Zn^2+^; TBD, TBD – apoferritin without Zn^2+^; TBD, TBD – aldolase. The 15, 30, and 60 eps datasets were deposited to the EMPIAR database under accession numbers TBD, TBD, and TBD respectively.

## Contributions

All authors contributed to the writing of the manuscript. BEB performed the coincidence loss experiments and the MTF, NPS, and DQE calculations using data collected on the FSU Titan Krios, RP prepared the apoferritin samples and collected the second apoferritin dataset, and the aldolase dataset on the Apollo. RP also performed the single particle analyses for those datasets and calculated all of the ResLog plots in the manuscript. XF collected the apoferritin datasets at the different dose rates and performed those single particle refinements. JHM collected the aldolase data on the K3 and the DE-64 and performed those single particle refinements. PSR did the apoferritin modeling. SS did the aldolase modeling.

